# QMaker: Fast and accurate method to estimate empirical models of protein evolution

**DOI:** 10.1101/2020.02.20.958819

**Authors:** Bui Quang Minh, Cuong Cao Dang, Le Sy Vinh, Robert Lanfear

## Abstract

Amino acid substitution models play a crucial role in phylogenetic analyses. Maximum likelihood (ML) methods have been proposed to estimate amino acid substitution models, however, they are typically complicated and slow. In this paper, we propose QMaker, a new ML method to estimate a general time-reversible *Q* matrix from a large protein dataset consisting of multiple sequence alignments. QMaker combines an efficient ML tree search algorithm, a model selection for handling the model heterogeneity among alignments, and the consideration of rate mixture models among sites. We provide QMaker as a user-friendly function in the IQ-TREE software package (http://www.iqtree.org) supporting the use of multiple CPU cores so that biologists can easily estimate amino acid substitution models from their own protein alignments. We used QMaker to estimate new empirical general amino acid substitution models from the current Pfam database as well as five clade-specific models for mammals, birds, insects, yeasts, and plants. Our results show that the new models considerably improve the fit between model and data and in some cases influence the inference of phylogenetic tree topologies.

## Introduction

Amino acid substitution models are crucial for model-based phylogenetic analyses of protein sequences, including for maximum likelihood (ML) and Bayesian inference approaches. Most commonly-used protein models are Markov processes summarised in a 20-by-20 replacement matrix, denoted as *Q*, which describes the rates of substitutions between pairs of amino acids. Because they have so many parameters, *Q* matrices are computationally very expensive to estimate. As a result, they are not usually estimated during a phylogenetic analysis of a single amino-acid multiple sequence alignment (MSA). Instead, the best *Q* matrix for each locus in a multilocus MSA is usually selected from a set of pre-estimated *Q* matrices using model selection software such as ModelFinder (Kalyaanamoorthy et al. 2017), ModelTest (Darriba et al. 2019), or PartitionFinder (Lanfear et al. 2017). Estimating *Q* matrices from large collections of empirical MSAs, where one derives the so-called *empirical Q* matrix that jointly explains substitution patterns across all MSAs, remains challenging both because the task is computationally expensive, and because there is no user-friendly software implementation that facilitates the task. As a result, the publication of new empirical *Q* matrices remains infrequent, and empirical phylogeneticists rarely estimate their own *Q* matrix even in those cases where they have sufficient data.

The first empirical *Q* matrices, Dayhoff (Dayhoff et al. 1978) and JTT (Jones et al. 1992), were estimated using the Maximum Parsimony (MP) principle. This approach simply counts the minimum number of amino-acid changes along a phylogeny required to explain the MSA. However, MP methods have a well-known shortcoming of not accounting for multiple amino-acid substitutions on single branches of the tree. Such shortcomings can be largely overcome by the Maximum Likelihood (ML) approach, where one estimates the *Q* matrix that maximises the joint likelihood of observing a large collection of MSAs given independently estimated tree topologies for each MSA. The most widely used *Q* matrices, WAG (Whelan and Goldman 2001) and LG (Le and Gascuel 2008), were estimated using the ML approach. These matrices substantially improved model fit on a range of MSAs compared with the older matrices. However, the methods used to estimate the LG and WAG matrices used several approximations to make the analyses computationally feasible. For example, Whelan and Goldman (2001) ignored rate heterogeneity across sites (RHAS), although this phenomenon is widely observed in empirical MSAs. Le and Gascuel (2008) later improved this method by incorporating RHAS with a discrete Gamma distribution (Yang 1994) but using a site-rate partition model instead of the originally designed mixture model. Moreover, the Pfam database (Bateman et al. 2002) used to estimate LG has now increased eight-fold (El-Gebali et al. 2019). As the most widely-used *Q* matrices were estimated more than a decade ago, improvements in the available data and phylogenetic inference methods suggest that it might be possible to estimate improved *Q* matrices.

### New Approaches

Here we present QMaker, an ML method and software implementation which estimates an empirical *Q* matrix for any set of protein MSAs. Figure 1 shows a schematic overview of the QMaker workflow (see Material and Methods for full details). QMaker improves upon previously published ML procedures on a number of fronts (Table 1). These include the use of the efficient ML tree search algorithm of IQ-TREE (Nguyen et al. 2015), consideration of a distribution-free model of RHAS (Kalyaanamoorthy et al. 2017), full usage of the rate mixture model, support for multiple CPU cores, and an explicit separation of training and testing data. Furthermore, we provide an easy-to-use implementation of QMaker as part of the IQ-TREE software package (http://www.iqtree.org). We employ our new software to estimate and compare seven new amino acid replacement matrices: two based on the most recent version of the Pfam database, and five clade-specific matrices for mammals, birds, insects, yeasts, and plants, respectively.

**Table 1:** Feature comparisons between QMaker and two previously published estimation procedures (Whelan and Goldman 2001; Le and Gascuel 2008).

| Feature | Whelan & Goldman | Le & Gascuel | QMaker |
| --- | --- | --- | --- |
| Tree reconstruction | Neighbor-joining (Saitou and Nei 1987) with Dayhoff+F distances | PhyML (Guindon and Gascuel 2003) with WAG+I+Γ4 model | IQ-TREE (Nguyen et al. 2015) with best-fit model |
| Branch length estimation | Scaled on JTT+F | ML estimate | ML estimate |
| Rate heterogeneity across sites | No | Gamma model (Yang 1994) | Gamma (Yang 1994), invariant site (Gu et al. 1995), and free rate model (Kalyaanamoorthy et al. 2017) |
| Optimisation technique | Expectation maximisation | Expectation maximisation | ML |
| Multi-core CPU support | No | No | Yes |
| Explicit separation of training and testing data | No | No | Yes |

**Figure 1.**
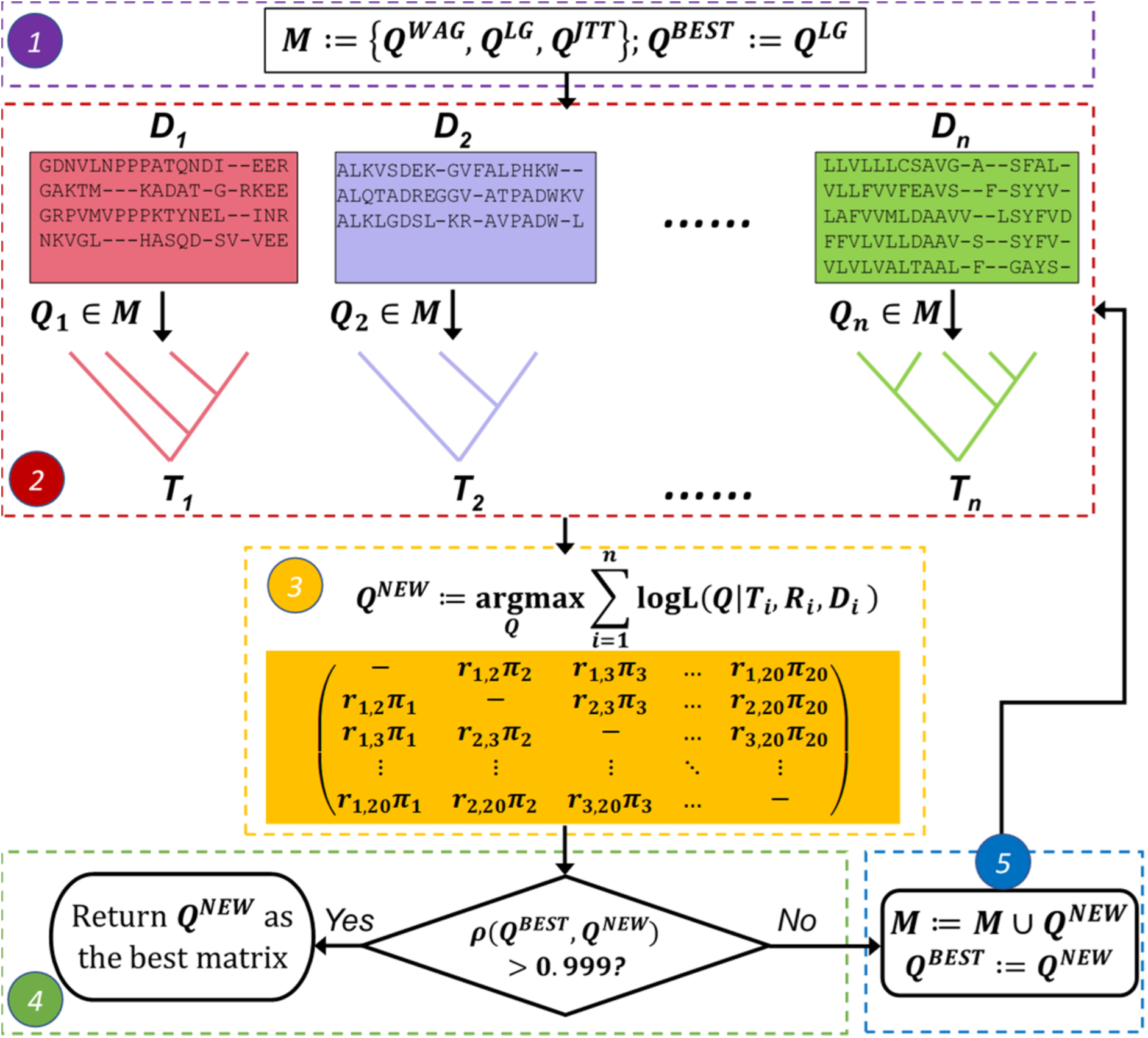
Schematic overview of QMaker consisting of five main steps: (1) initialize the current best replacement matrix *Q*^*BEST*^ as LG and the set of candidate matrices as WAG, LG, and JTT; (2) for each alignment_*D*_*i*_ find the best-fit matrix *Q*_*i*_ and the best ML tree *T*_*i*_; (3) maximize the joint log-likelihood to obtain a new matrix *Q*^*NEW*^ that best explains all *D*_*i*_; (4) if the Pearson correlation between *Q*^*BEST*^ and *Q*^*NEW*^ is higher than 0.999, return *Q*^*NEW*^ as the best matrix for the data; (5) otherwise, replace *Q*^*BEST*^ by *Q*^*NEW*^, extend the set of candidate matrices with *Q*^*NEW*^ and go back to step 2.

## Results

### QMaker outperforms existing estimation methods

To establish whether the approach we propose here improves upon previously-suggested approaches, we compared QMaker (Figure 1; Materials and Methods) to the method of Le and Gascuel (2008) on the same training data. Because both methods use the same data to estimate a *Q* matrix, any differences between the matrices must be due solely to the estimation procedure. Such differences can then be assessed using information theoretic approaches.

To compare the two approaches, we first downloaded the training set of 3,412 Pfam MSAs originally used to estimate the LG matrix and then applied QMaker to estimate a new *Q* matrix from this data. We called the resulting matrix Q.LG to reflect the origin of the dataset. QMaker took about 28 hours wall-clock time using 36 CPU cores on a 2.3-GHz server to complete the whole training.

To compare the performance of the LG and Q.LG matrix, we asked how frequently each matrix was selected as the best-fit model for the 500 test MSAs originally used to test the LG matrix (Le and Gascuel 2008). To do this, we calculated the best-fit model for each MSA from a set of four candidate *Q* matrices comprised of the two comparator matrices LG and Q.LG, plus two other frequently-used matrices: WAG (Whelan and Goldman 2001) and JTT (Jones et al. 1992). Q.LG was the most frequently selected matrix (232 MSAs), followed by LG (136 MSAs), WAG (61 MSAs), and JTT (44 MSAs). These results demonstrate that the QMaker method improves the fit between models and data compared to previous estimation procedures. For the sake of reproducibility, we provide the Q.LG matrix in the supplementary material. We do not, however, intend for the Q.LG matrix to be widely used, as it is estimated from a now-outdated version of the Pfam database.

### Larger amino acid databases improve model fit, but primarily to target alignments

We used QMaker to estimate two new amino-acid substitution matrices from the latest version of the Pfam database: Q.pfam from a training set of 6,654 MSAs of the Pfam database version 31 (El-Gebali et al. 2019); and Q.pfam-gb from 3,742 training MSAs of the same database, but for which the MSAs were pre-processed with GBlocks (Castresana 2000) to remove gappy and non-conserved sites, and to remove MSAs with low numbers of sites and/or sequences (Materials and Methods). We then compared the fit of Q.pfam and Q.pfam-gb to three previously-estimated matrices (LG, WAG, and JTT) using two sets of test MSAs. The first test set comprises 6,654 Pfam MSAs that were not used to estimate either of the new matrices. The second set is a subset of the first, and comprises the 3,727 MSAs that remain after applying GBlocks and filtering out MSAs with low numbers of sites and/or sequences from the 6,654 MSA test set (Materials and Methods).

Q.pfam and Q.pfam-gb outperformed other matrices on the test MSAs, with each matrix being the best fit to the data corresponding to that on which it was trained. Q.pfam was the most frequently selected matrix on first test set (34.2%), followed by LG (26.7%), Q.pfam-gb (15.9%), JTT (14.1%), and WAG (9.1%). Similarly, Q.pfam-gb was the most frequently selected matrix for the second test set (38.4%), followed by LG (25.5%), JTT (16.1%), Q.pfam (14.0%), and WAG (6.0%).

We further tested the new matrices on a collection of 13,041 single-locus MSAs from five recently-published phylogenomic datasets (Table 2). To do this, we compared the fit of the same five models (Q.pfam, Q.pfam-gb, LG, WAG, and JTT) to each of the 13,041 MSAs. Surprisingly, the most commonly-selected matrix across all 13,041 MSAs was JTT (74.9%), followed by Q.pfam-gb (14.3%), LG (5.9%), Q.pfam (3.0%), and WAG (2.0%). The JTT matrix was the most commonly-selected matrix for three out of the five datasets (Birds, Plants, and Mammals; supplementary figure S1), and the Q.pfam-gb matrix was the most commonly-selected matrix for the remaining two datasets (Insects and Yeasts; supplementary figure S1). This shows that amino acid models estimated from the Pfam database (Q.pfam, Q.pfam-gb, LG) often fail to provide the best fit to alignments used for phylogenomic inference on commonly-studied clades.

**Table 2:** Summary of the datasets used to estimate new amino-acid replacement matrices. For each dataset we randomly subsampled half (Pfam) or 1,000 MSAs (others) as the training set and remaining loci as the test set. For bird and plant datasets we used two non-overlapping training sets to examine the effect of random subsampling. For the yeast dataset, we additionally subsampled 100 sequences from the training set due to the excessive computational burden.

| Dataset | Reference | Seqs | Sites | Loci | Training | Testing |
| --- | --- | --- | --- | --- | --- | --- |
| Pfam | (El-Gebali et al. 2019) | 1,150,099 | 3,433,343 | 13,308 | 6,654 | 6,654 |
| Pfam-gblocks | (El-Gebali et al. 2019) | 420,433 | 1,032,972 | 7,469 | 3,742 | 3,727 |
| Plant | (Ran et al. 2018) | 38 | 432,014 | 1,308 | 1,000 | 308 |
| Bird | (Jarvis et al. 2015) | 52 | 4,519,041 | 8,295 | 1,000 × 2 | 6,295 |
| Mammal | (Wu et al. 2018) | 90 | 3,050,199 | 5,162 | 1,000 × 2 | 3,162 |
| Insect | (Misof et al. 2014) | 144 | 595,033 | 2,868 | 1,000 | 1,868 |
| Yeast | (Shen et al. 2018) | 343 | 1,162,805 | 2,408 | 1,000<br>*100 seqs* | 1,408 |

### Five new clade-specific Q matrices improve model fit on phylogenomic data

The surprisingly poor fit of Pfam-based matrices (Q.pfam, Q.pfam-gb, LG) to the empirical MSAs, combined with the high variation in the identity of the best model in each dataset, suggests that there may be substantial between-clade variation in the way that proteins used for phylogenetic inference evolve. If this is the case, then accounting for this by estimating independent *Q* matrices for each clade should improve model fit. To test this, we estimated a clade-specific *Q* matrix for each of the five phylogenomic datasets (Table 2): Q.plant, Q.bird, Q.mammal, Q.insect, and Q.yeast. For each dataset we used 1000 training MSAs to estimate the *Q* matrix, and the remaining MSAs from each dataset as test sets (see Materials and Methods for more details).

Figure 2 shows the frequency with which each of the seven new (Q.pfam, Q.pfam-gb, Q.plant, Q.bird, Q.mammal, Q.insect, and Q.yeast) and three existing (JTT, WAG, and LG) matrices were selected as the best-fit for the seven test sets. As expected, the best fit *Q* matrix for each test set was the *Q* matrix estimated from the corresponding training set, although the strength of the association varied widely among test datasets. For example, Q.plant was the best model for 90.1% of plant test MSAs, with the next best model selected for fewer than 5% of test MSAs. But Q.pfam-gb was only selected as the best model for 25.2% of the Pfam-gb test MSAs, with the next best model (LG) selected for almost 20% of MSAs. These results are likely driven in part by the fact that the set of models we considered included many models that are similar to Q.pfam-gb (e.g. Q.pfam, LG, WAG), but few that are similar to Q.plant.

**Figure 2.**
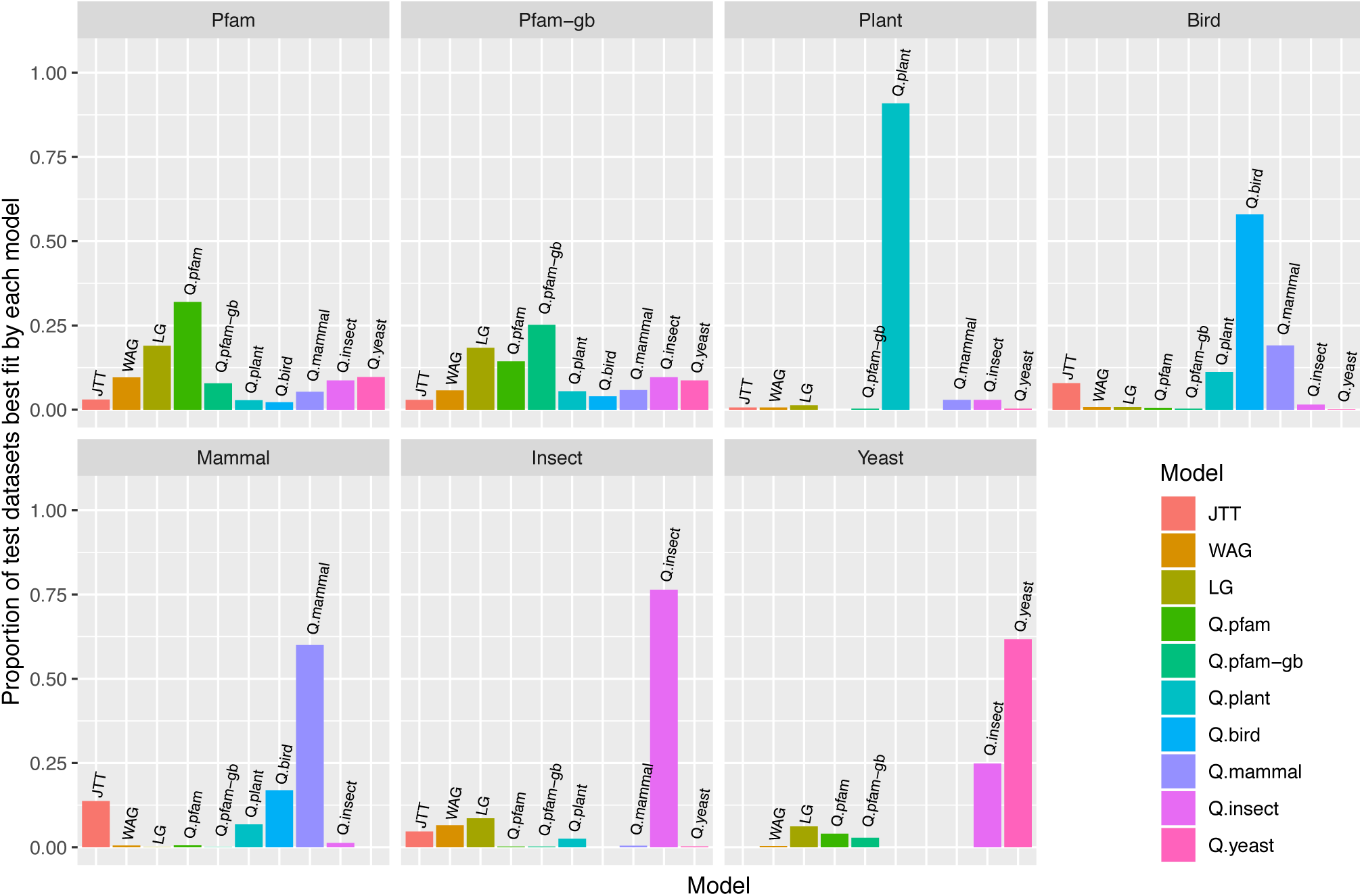
Frequency of best fitting for 10 amino-acid replacement matrices on 7 test datasets: Pfam, Pfam processed with GBlocks, Plant (Ran et al. 2018), Bird (Jarvis et al. 2015), Mammal (Wu et al. 2018), Insect (Misof et al. 2014), and Yeast (Shen et al. 2018).

### Principle components analyses reveal the landscape of amino acid models

We used principle components analyses (PCA) to compare the properties of the seven new amino acid models presented here to 19 previously-estimated models (Table 3). The PCA plot of the *Q* matrices (Figure 3A) shows a clear separation between matrices inferred from the nuclear, mitochondrial, chloroplast and viral genomes and the clade-specific matrices, with the clade-specific matrices falling between the mitochondrial and viral matrices, and the three Pfam-based matrices (LG, Q.pfam and Q.pfam-gb) in close proximity. The PCA plot of the models’ amino acid frequencies (Figure 3B) reveals that most of the variation among frequency vectors comes from differences between and within the viral and mitochondrial models, with more limited separation between the clade-specific matrices (Q.bird, Q.plant, Q.insect, Q.mammal, Q.yeast) and the general-purpose matrices (LG, WAG, JTT, Q.pfam, Q.pfam-gb).

**Table 3:**
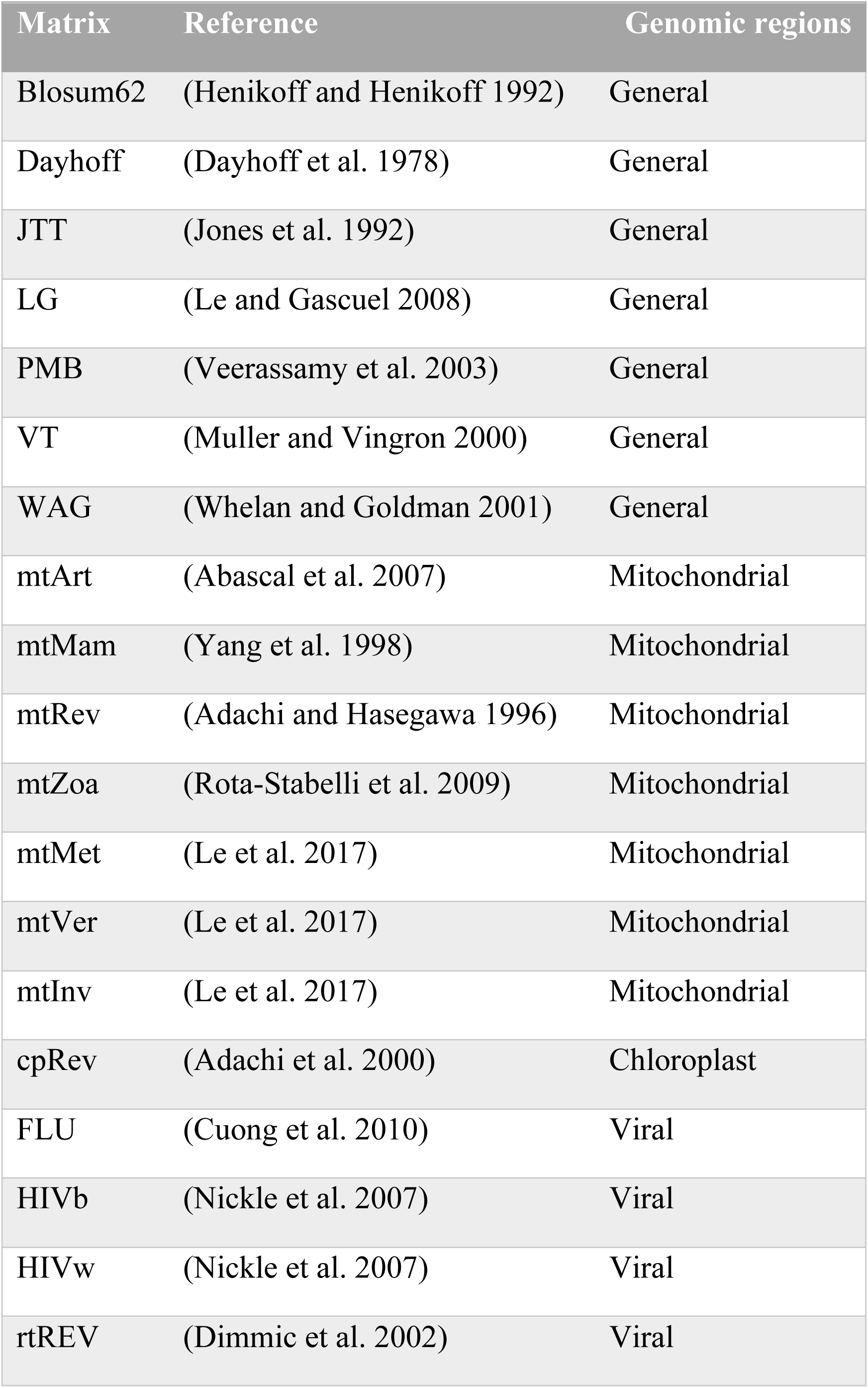
Existing amino-acid replacement matrices.

| Matrix | Reference | Genomic regions |
| --- | --- | --- |
| Blosum62 | (Henikoff and Henikoff 1992) | General |
| Dayhoff | (Dayhoff et al. 1978) | General |
| JTT | (Jones et al. 1992) | General |
| LG | (Le and Gascuel 2008) | General |
| PMB | (Veerassamy et al. 2003) | General |
| VT | (Muller and Vingron 2000) | General |
| WAG | (Whelan and Goldman 2001) | General |
| mtArt | (Abascal et al. 2007) | Mitochondrial |
| mtMam | (Yang et al. 1998) | Mitochondrial |
| mtRev | (Adachi and Hasegawa 1996) | Mitochondrial |
| mtZoa | (Rota-Stabelli et al. 2009) | Mitochondrial |
| mtMet | (Le et al. 2017) | Mitochondrial |
| mtVer | (Le et al. 2017) | Mitochondrial |
| mtInv | (Le et al. 2017) | Mitochondrial |
| cpRev | (Adachi et al. 2000) | Chloroplast |
| FLU | (Cuong et al. 2010) | Viral |
| HIVb | (Nickle et al. 2007) | Viral |
| HIVw | (Nickle et al. 2007) | Viral |
| rtREV | (Dimmic et al. 2002) | Viral |

**Figure 3.**
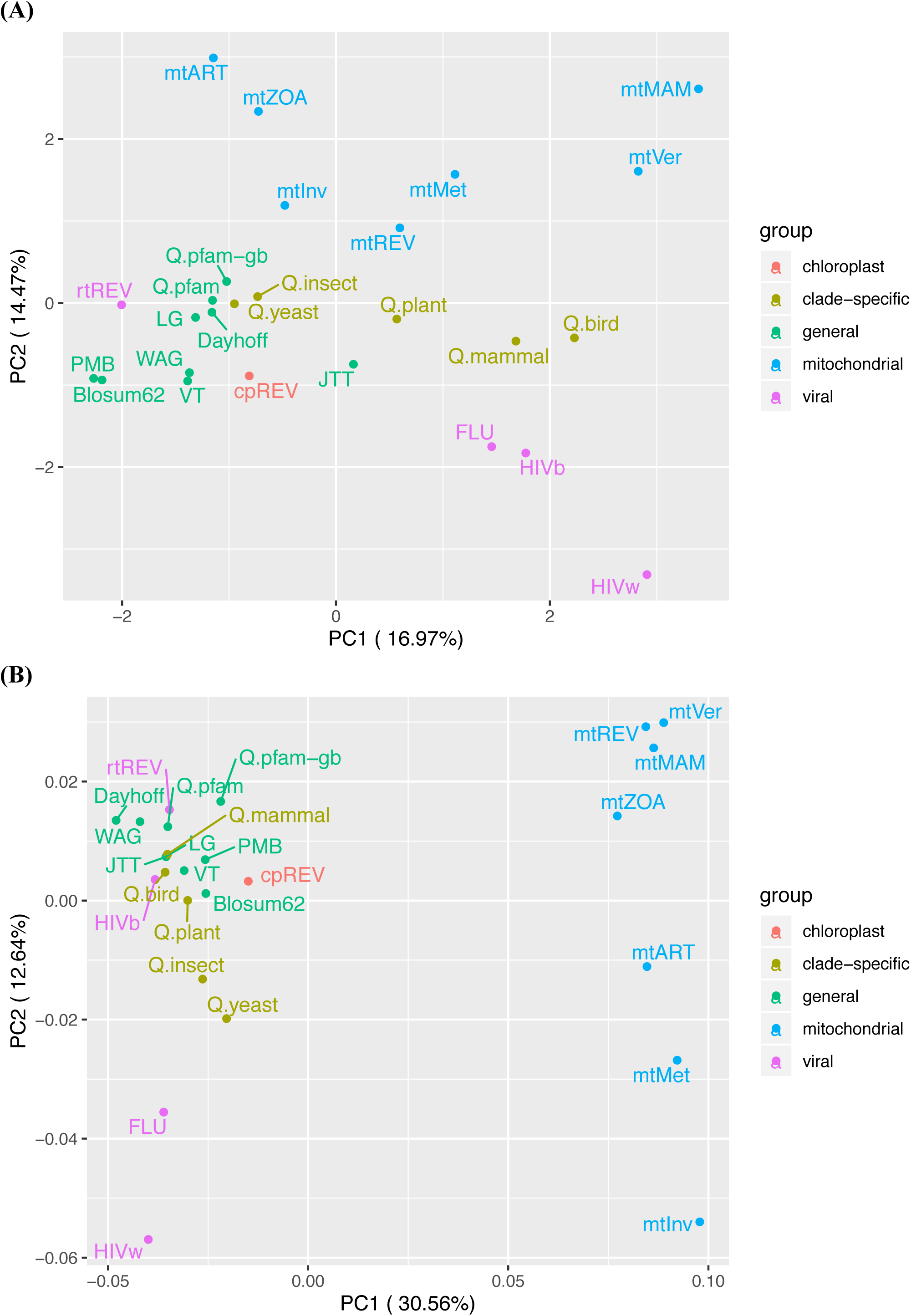
Principle component analysis (PCA) of all matrices with respect to (A) the amino acid exchangeability rates and (B) amino-acid frequencies.

### Incorporating the new matrices into model selection changes locus-tree inference

To examine whether the seven new matrices we propose here affect the inference of phylogenetic trees, we asked how often the tree changed when one of the new models was selected as the best model. For each single-locus MSA in each dataset, if one of the new models was selected as best model, we inferred the ML locus tree using the new model which we denoted *T*_*new*_. We then compared this tree to the tree inferred for the same MSA using the best-fit model among JTT, WAG and LG, which we denoted *T*_*old*_. Differences between *T*_*new*_ and *T*_*old*_ could come from two sources: the effects of using a different amino acid substitution model or the stochasticity in tree search. To decouple these two factors we then performed another independent tree search to infer *T*_*old*2_ in the same say as inferring *T*_*old*_ but using a different random number seed. If *T*_*old*_ is different from *T*_*old*2_, then the difference is merely due to tree search stochasticity. For each dataset, we then compared the distribution of normalized Robinson-Foulds (nRF) (Robinson and Foulds 1981) distances between *T*_*new*_ and *T*_*old*_ to the distribution of the nRF distances between *T*_*old*_ and *T*_*old*2_. The extent to which nRF distances between *T*_*new*_ and *T*_*old*_ are larger than those between *T*_*old*_ and *T*_*old*2_ indicates the extent to which the new model affects tree inference, independently of stochasticity in the tree search.

Figure 4 shows the distributions of nRF(*T*_*new*_, *T*_*old*_) and nRF(*T*_*old*_, *T*_*old*2_) for the seven test datasets. Figure 4 shows noticeably higher nRF distances between *T*_*new*_ and *T*_*old*_ compared to *T*_*old*_ and *T*_*old*2_, indicating that using the newly proposed (and better fit) models changes locus tree topologies in every dataset. In fact, the two distributions are significantly different for all datasets (p < 0.001 from a Kolmogorov-Smirnov test comparing the two distributions in each dataset), indicating that the new models of evolution affect a non-trivial number of single-locus tree topologies in every dataset.

**Figure 4.**
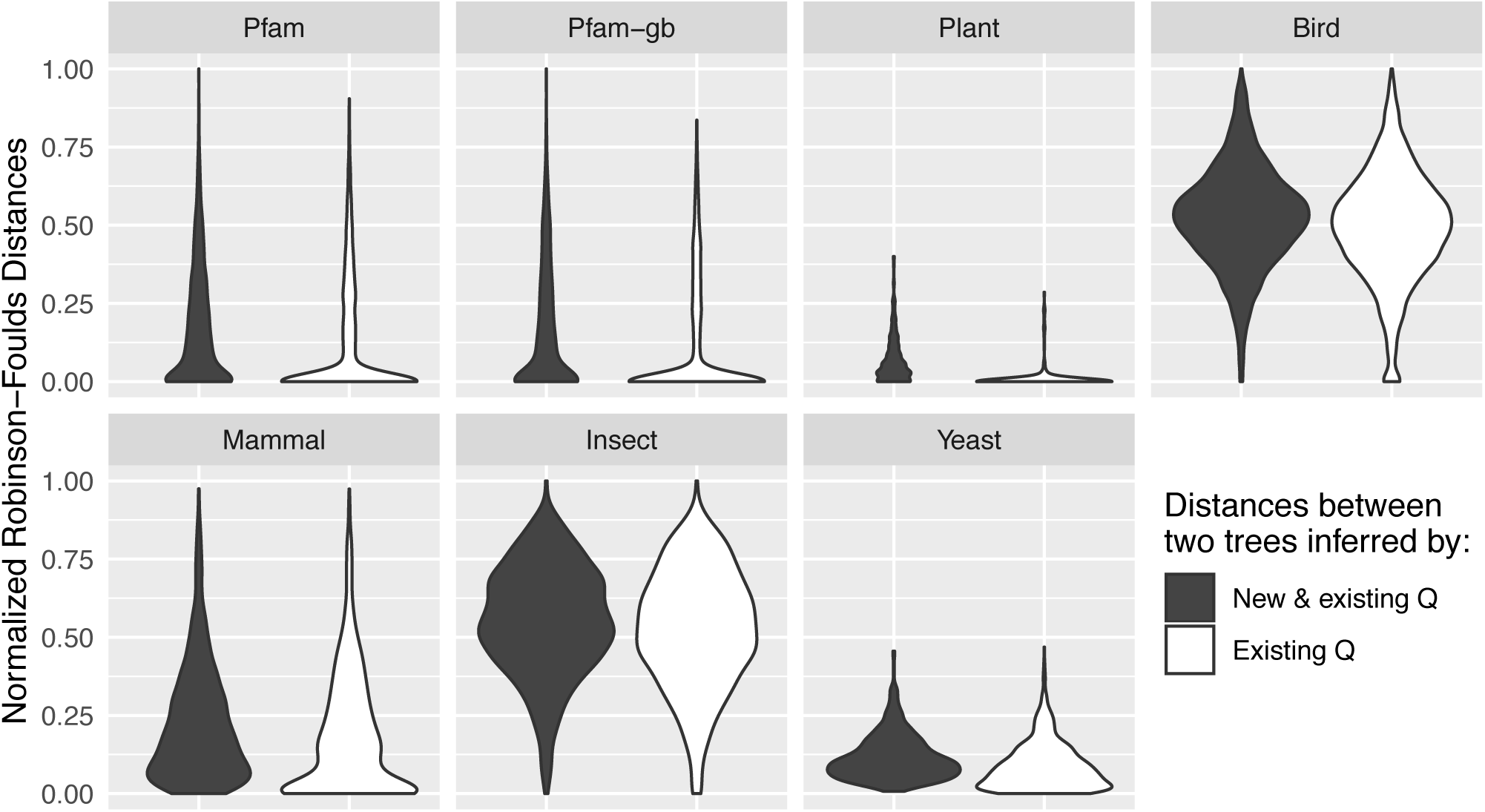
Normalized Robinson-Foulds (nRF) distances between the gene trees inferred by new and existing (JTT, WAG, or LG) models.

## Discussions

In this study, we describe and implement QMaker, an easy-to-use tool to estimate an amino acid replacement matrix Q for any dataset of one or more amino-acid alignments. Phylogenetic inference from amino acid alignments relies heavily on pre-computed *Q* matrices. It is a little surprising, therefore, that new *Q* matrices are published relatively infrequently (e.g. Table 3), particularly in the age of phylogenomics when an increasing number of studies collect sufficient data to reliably estimate a *Q* matrix. We hope that the development of QMaker will democratise the inference *Q* matrices, and that this will lead to concomitant improvements in phylogenetic inference and our understanding of molecular evolution.

The approach we implement in QMaker builds on previously-described approaches (Whelan and Goldman 2001; Le and Gascuel 2008), and our analyses reveal that it improves on them in terms of model fit to the data. We applied QMaker to estimate two general-purpose *Q* matrices and five clade-specific *Q* matrices (mammals, plants, birds, insects and yeasts). We showed that they not only improve the fit between the model and the data but also influence the tree topologies. All of the new matrices are now implemented in IQ-TREE version 2 (Minh et al. 2020) and incorporated as part of the model selection procedure, and the data necessary to implement all matrices in other phylogenetic software packages are provided in the supplementary material. We hope that the addition of these matrices will improve phylogenetic inference for researchers who do not have sufficient data to estimate a *Q* matrix.

The relationships among the 19 existing *Q* matrices and the 7 new matrices we present here (Figure 3) reveal a number of interesting patterns. As expected, there is a clear distinction between *Q* matrices estimated from different genomes, with the general purpose matrices estimated from large datasets of protein alignments from the nuclear genome tending to cluster tightly together (Figure 3). More surprising is the observation that the five new clade-specific *Q* matrices we estimate here tend to be quite distinct from all other *Q* matrices, and are also remarkably distinct from one another. This result, combined with our observations that the clade-specific *Q* matrices tend to improve model fit and affect tree inference, highlight the potential benefits of inferring a clade-specific *Q* matrix before inferring a phylogeny. The differences among the clade-specific matrices also hint at potentially significant differences between the molecular evolutionary processes driving protein evolution in different clades of organisms.

The sometimes substantial variation in best-fit model for different loci from a single dataset (Figure 2) confirms that there can also be substantial variation in molecular evolution among loci. Thus, although QMaker allows researchers to infer a single *Q* matrix from a collection of alignments, it still seems sensible to infer phylogenies in a framework that allows for different *Q* matrices to be applied to different loci, such as by using partitioned (Lanfear et al. 2012; Chernomor et al. 2015) or mixture models.

The QMaker framework opens new avenues of research by simplifying the process of inferring a single *Q* matrix, but is currently limited to estimating a single reversible *Q* matrix from a one or more amino acid alignments. In principle, both of these limitations could be relaxed, for example by extending the QMaker approach to infer non-reversible *Q* matrices (e.g., Minh et al. 2020) and/or mixtures of *Q* matrices from amino acid alignments (e.g. as was done to infer the LG4M and LG4X mixtures of matrices (Le et al. 2012). Both of these approaches have the potential to further improve phylogenetic inference beyond the developments that we present here.

## Material and Methods

### Datasets used for training and testing

We downloaded a total of 16,712 Pfam MSAs from version 31 of the database (El-Gebali et al. 2019), removed identical sequences from each MSA and only retained MSAs having between 5 and 1,000 sequences and at least 50 sites. This leaves us with 13,308 remaining MSAs, denoted as the Pfam dataset. We also applied GBlocks (Castresana 2000) to filter out potentially mis-aligned sites (e.g. due to high sequence divergence), which we call Pfam-gb. Moreover, we downloaded five datasets for Plant (Ran et al. 2018), Bird (Jarvis et al. 2015), Mammal (Wu et al. 2018), Insect (Misof et al. 2014), and yeast (Shen et al. 2018). For each of the seven datasets we divided the loci into two subsets: a training set to estimate Q and a test set to compare the model fit between the estimated Q. Details of the datasets are summarized in Table 2. All data are available from the supplementary material and https://github.com/roblanf/BenchmarkAlignments (clade-specific datasets).

### Model of amino acid substitutions

The model of amino acid substitutions follows the continuous Markov process that is stationary, reversible and homogeneous. This process is summarised in a 20-by-20 rate matrix *Q*, that describes the rate of change from one amino acid to another per time unit. Because of the reversibility assumption, entries of *Q* can be written as the product of the symmetric exchangeability rates (*r*_*ij*_) and the amino-acid frequencies (*π*_*i*_):

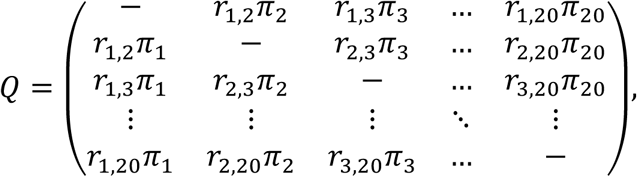

where the diagonal entries of *Q* are chosen such that its row sums equal 0.

The 190 exchangeability parameters *r*_*ij*_ can only be reliably estimated from a large amount of data. Therefore, almost all phylogenetic analyses on protein sequences use a pre-determined (*r*_*ij*_) matrix such as WAG (Whelan and Goldman 2001) and LG (Le and Gascuel 2008). These matrices were obtained by a survey of large protein databases such as BRKALN and Pfam. Quite often the amino acid frequencies can be empirically estimated from the dataset at hand, denoted by “+F”. For example, the WAG+F model uses the exchangeability rates defined by WAG but amino acid frequencies from the current MSA.

### Model of rate heterogeneity across sites

It is well known that MSA sites may have evolved at different rates. The so-called rate heterogeneity across sites (RHAS) has been typically modelled by a discrete Gamma distribution (Yang 1994) w/o a proportion of invariable sites (Gu et al. 1995). For example, LG+I+Γ means that while all sites follow the LG exchangeability matrix, a fraction of sites is invariable (i.e. with zero evolutionary rates due to e.g. selective pressure) and the rates of the remaining variable sites follow a Gamma distribution.

Recently, it has been shown that the assumption of Gamma distribution is not justifiable and a distribution-free rate model frequently provides a better fit (Kalyaanamoorthy et al. 2017). Such model is denoted by e.g., LG+R5, that means sites fall into five rate categories with no distributional constraints on the rates and proportions of each category, which will be estimated by ML.

### Estimating a joint replacement matrix from a protein dataset

We now introduce a new method to estimate a replacement matrix *Q* from a database of protein MSAs *D =* {*D*_1_, …, *D*_*n*_}. Here, we want to find a single *Q* that best explains, in terms of maximum likelihood, the pattern of amino acid substitutions for all MSAs. We denote by *M =* {*Q*^*WAG*^, *Q*^*LG*^, …} the set of candidate replacement matrices (Table 3).

We first determine, for each *D*_*i*_, the best-fit matrix *Q*_*i*_ ∈ *M*, the best RHAS model *R*_*i*_ (e.g. I+Γ and R5) using ModelFinder (Kalyaanamoorthy et al. 2017), and the ML tree *T*_*i*_ with the set of branch lengths Λ_*i*_ using IQ-TREE (Nguyen et al. 2015). Next, we fix *R*_*i*_, *T*_*i*_ and Λ_*i*_ to estimate the *Q* that maximizes the total log-likelihood across all MSAs in *D*:

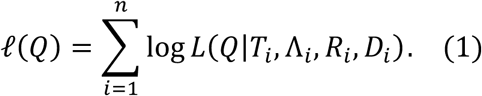

This maximisation results in an estimate of a new rate matrix denoted as *Q*^*NEW*^. Some MSAs may now show *Q*^*NEW*^ as the better-fit model. Therefore, we extend the set of candidate rate matrices by *Q*^*NEW*^ and repeat the procedure above to re-estimate *Q*_*i*_, *R*_*i*_, *T*_*i*_, Λ_*i*_. The overall workflow of QMaker is as follows (Figure 1):

1. Initialise the set of candidate replacement matrices as *M* := {*Q*^*WAG*^, *Q*^*LG*^, *Q*^*JTT*^} and the current best matrix as *Q*^*BEST*^ := *Q*^*LG*^.
2. For each MSA *D*_*i*_, find the best-fit matrix *Q*_*i*_ ∈ *M*, rate heterogeneity across sites model *R*_*i*_ and estimate ML tree *T*_*i*_ with branch lengths Λ_*i*_ based on *Q*_*i*_ and *R*_*i*_.
3. Given *R*_*i*_ and *T*_*i*_, jointly estimate *Q* and Λ_*i*_ that maximises the log-likelihood function (1), resulting in a new replacement matrix *Q*^*NEW*^. Specifically, 3a) Let *k* := 0 be the number of jointly estimated rounds 3b) estimate *Q* given *R*_*i*_, *T*_*i*_, and Λ_*i*_ 3c) estimate Λ_*i*_ given *R*_*i*_, *T*_*i*_, and *Q;* 3*d*) increase *k* := *k* + 1. If *k* is smaller than a predefined threshold, go to step 3b, otherwise, go to step 4.
4. Let *ρ* be the Pearson correlation between *Q*^*BEST*^ and *Q*^*NEW*^. If *ρ* > 0.999, report *Q*^*NEW*^ as the best replacement matrix for the database *D* and stop. Otherwise, go to step 5.
5. Assign *Q*^*BEST*^ := *Q*^*NEW*^, add *Q*^*NEW*^ to the set of candidate matrices: *M* := *M* ∪ *Q*^*NEW*^, and go back to step 2.

### Comparisons with previous estimation procedures

Compared with the ML procedures used to estimate WAG (Whelan and Goldman 2001) and LG (Le and Gascuel 2008), QMaker has a number of differences (Table 1). Among others, Whelan and Goldman (2001) omitted rate heterogeneity across sites and employed neighbour-joining for computational efficiency. Le and Gascuel (2008) improved this method by incorporating the Γ model of rate heterogeneity and inferring the ML tree with PhyML. However, they did not use the original Γ as a mixture model when estimating *Q*. Rather, they partitioned the sites in each *D*_*i*_ into their most likely rate category resulting in 4 sub-MSAs for each *D*_*i*_, and essentially applied the method of Whelan and Goldman to derive the *Q* matrix from the expanded *D*.

Here, QMaker improves both methods by (i) additionally considering the free rate and invariant site mixture models; (ii) inferring the ML tree with IQ-TREE, which has been shown to outperform PhyML and other software (Zhou et al. 2018); and (iii) directly optimising the log-likelihood function (1) to obtain Q instead of aforementioned approximations.

### Software implementation

We provided an implementation of QMaker as part of the IQ-TREE software. The entire training stage for the Pfam dataset can be accomplished with just two command lines. The first one is

~~~
iqtree -S ALN_DIR -nt 48
~~~

to find the best-fit models and ML trees for all MSAs residing in the folder `ALN_DIR`; `-nt` option is to specify the number of CPU cores. Note that for this study, due to the excessive size of the Pfam training set, we additionally used two options: `-mset LG,WAG,JTT` to consider only these three models and -`cmax 4` to restrict up to four categories for the rate heterogeneity across sites model. The second command line is

~~~
iqtree -S ALN_DIR.best_model.nex -nt 48 -te ALN_DIR.treefile -
-model-joint GTR20+FO
~~~

to perform step 3 of estimating the replacement matrix (GTR20 for general time reversible model with 20-state data), given the best models (`ALN_DIR.best_model.nex`) and best trees (`ALN_DIR.treefile`) found above.

For the five clade-specific datasets we performed an edge-linked partition model, which assumes a single tree topology and rescales edge lengths across the loci. This model was shown to best balance between model parameterization (Duchene et al. 2019). For this purpose, the `-S` option is changed to `-p`.

To test the model fit of the trained *Q* matrices we ran ModelFinder (Kalyaanamoorthy et al. 2017) as implemented in IQ-TREE:

~~~
iqtree -S TEST_DIR -m MF -mset JTT,WAG,LG,Q.pfam,Q.pfam-
gb,Q.plant,Q.bird,Q.mammal,Q.insect,Q.yeast
~~~

where `TEST_DIR` is a directory containing the testing MSAs.

## Supplementary material

Supplementary materials are available from https://doi.org/10.6084/m9.figshare.9768101.

## Acknowledgements

This research is funded by Vietnam National Foundation for Science and Technology Development (NAFOSTED) under grant number 102.01.2019.06.

